# Heparan Sulfate Controls Nanoscale Assembly of GPC3-Wnt Receptor Complexes

**DOI:** 10.64898/2026.07.21.739947

**Authors:** Shaoli Lin, David A. Ball, Mohamadreza Fazel, Tatiana S. Karpova, Mitchell Ho

## Abstract

Glypican-3 (GPC3) is a heparan sulfate proteoglycan that is highly expressed in hepatocellular carcinoma and promotes tumor progression through Wnt3a/β-catenin signaling. However, how the nanoscale organization of GPC3 at the cell surface controls signaling remains unclear. Here, we combined nano-resolution MINFLUX imaging, single-molecule tracking, and functional assays to define the spatial architecture and dynamics of GPC3 on hepatoma cells. We found that GPC3 exists as both single molecules and nanoscale clusters and switches between confined and free diffusions on the plasma membrane. Heparan sulfate (HS) chains create nanoscale corrals that limit GPC3 movement, whereas removal of HS increases diffusive heterogeneity and disrupts confinement. Wnt3a stimulation induces the formation of higher-order GPC3 assemblies and enhances β-catenin signaling, while loss of HS markedly reduces this response. MINFLUX DNA-PAINT further revealed that HS chains orchestrate the spatial distribution of Wnt3a and promote its association with the Wnt receptor, Frizzled-1, an essential step for pathway activation. Collectively, these findings reveal that HS controls the nanoscale organization and dynamics of GPC3 to promote Wnt receptor assembly and efficient β-catenin signaling in hepatoma cells.

## Introduction

Primary liver cancer (PLC) has a high mortality rate with increasing incidence rate in the United States, resulting in nearly a million new cases worldwide annually and making it one of the leading causes of death due to cancer ^1^. Hepatocellular carcinoma (HCC) is a major form of primary liver cancer, accounting for nearly all cases ^2^. Glypican-3 (GPC3) is a heparan sulfate proteoglycan (HSPG), which is prominently expressed on the surface of HCC cells and has emerged as a focal point in the pursuit of therapeutic interventions ^3,4^. GPC3 exists in four isoforms, with isoform 2 being the most prevalent, encoding a 70 kDa core protein ^5^. This full-length GPC3 protein undergoes furin-mediated processing, resulting in a 40 kDa N-terminal protein, that is released into the serum, and a 30 kDa C-terminal membrane-bound protein that is tethered to the cell membrane via a glycosylphosphatidylinositol (GPI) anchor. Two heparan sulfate chains are linked to the C-terminus of GPC3 through amino acids S495 and S509 ^5,6^.

High expression of GPC3 is strongly correlated with tumor progression and it promotes tumor growth through activating Wnt3a signaling in liver cancer cells ^7^. Briefly, the Wnt3a molecule acts through both the heparan sulfate (HS) chains and core protein of GPC3 to initiate the Wnt3a/β-Catenin signaling for tumor progression ^8,9^, presumably by forming a complex with the transmembrane Frizzled (FZD) cellular receptor family. The FZD family and the associated adaptors then transduce the downstream signals and promote the translocation of β-Catenin into the nucleus for cell survival gene transcription in the cells. Blocking of the Wnt3a binding site of GPC3 with antibodies results in the impairment of Wnt3a/β-Catenin signaling and tumor migration ^9,10^. Given the importance of Wnt3a/GPC3/β-Catenin signaling in liver cancer cells, understanding the spatial distribution and dynamics of GPC3 on the cell surface may shed light on the signaling modulation and the potential targeting strategy of GPC3. Therefore, in this study, we aimed to characterize the spatial organization and dynamic behavior of GPC3 to determine how its biophysical properties relate to its biological function.

We used the cutting-edge tools of single molecule localization (Nanoscopy) and Single Molecule Tracking (SMT) to assay dynamics of membrane-bound GPC3 in hepatoma cells in culture ^11–13^. We showed that GPC3 was concentrated and distributed along the adhesion structure on the cell surface. Quantitative trajectory analysis demonstrated that clustered GPC3 exhibits restricted subdiffusive motion, whereas dispersed molecules undergo near-Brownian diffusion. HS chains promote GPC3 assembly and stabilize nanoscale corralling, and Wnt3a stimulation induced spatial reorganization of GPC3 consistent with signaling complex assembly. Together, these findings establish a direct relationship between the nanoscale architecture of GPC3, its diffusion behavior, and its signaling competence, linking membrane organization to Wnt3a pathway activation in cancer cells.

## Results

### GPC3 associates with ER-Golgi network in the cytoplasm and resides on lipid rafts on the cell membrane

Little is known about the distribution of GPC3 in hepatoma cells. To explore the GPC3 distribution, we expressed mCherry-GPC3 in Hep3B-GPC3-KO (SC22) hepatoma cells and found that GPC3 localizes both in the cytoplasm and on the cytoplasmic membrane (Fig. 1A). The HS chains are synthesized and attached to glycoproteins in the Golgi apparatus of cells. This process involves many enzymes that can modify the structure of the chains ^14^, therefore we proposed that the intracellular GPC3 localized to the Golgi. To test this hypothesis, we expressed GPC3 in Hep3B-SC22 cells together with an ER or Golgi marker (Calnexin and GalT, respectively). Confocal microscopy shows that GPC3 co-localized exclusively with GalT and partially with Calnexin, indicating GPC3 is preferentially located on Golgi in the cytoplasm (Fig. 1B).

**Figure 1.**
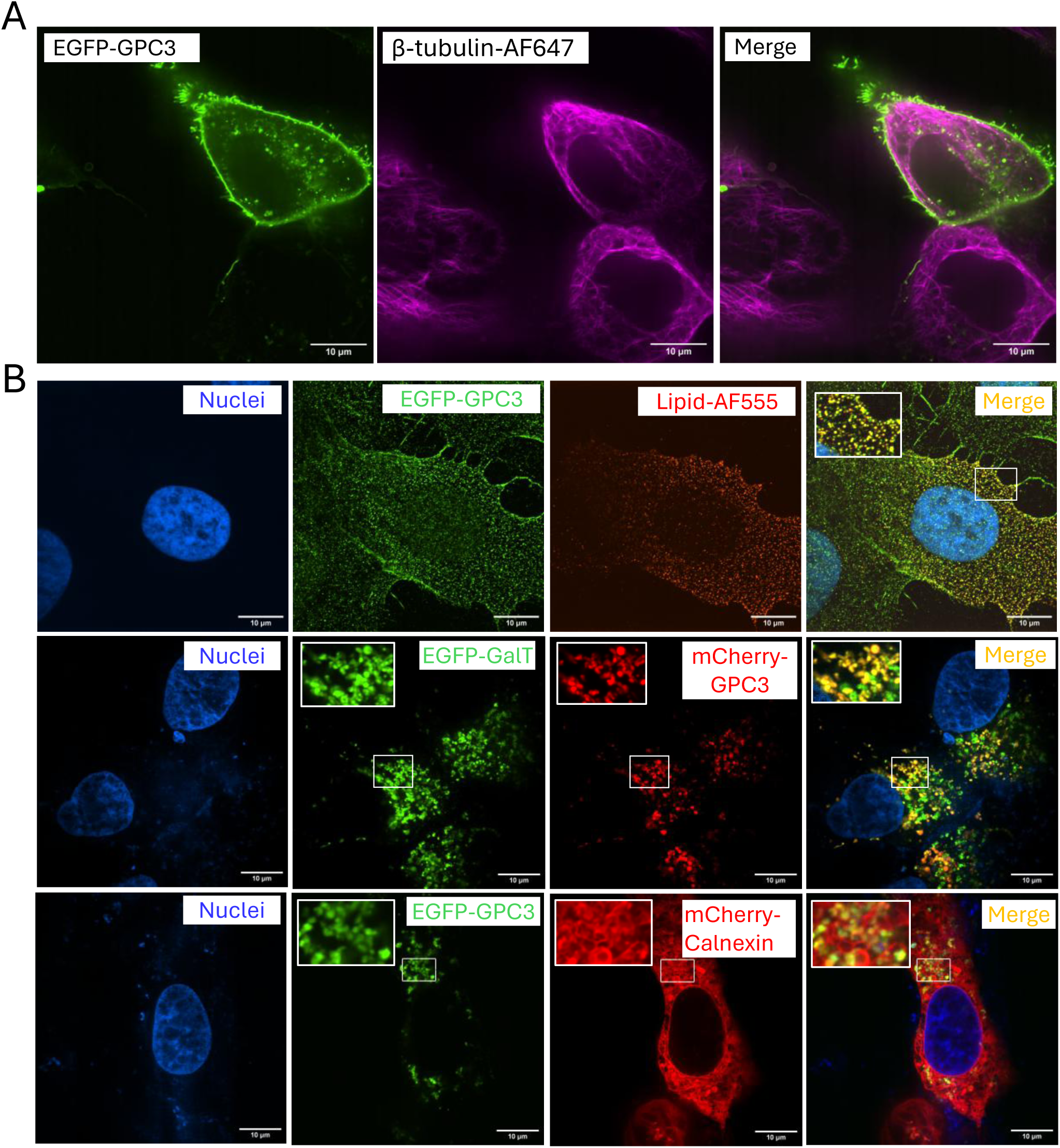
Co-localization of GPC3 and lipid on cell membrane and Golgi in the cytoplasm. A. Membrane and cytoplasmic distribution of GPC3. EGFP-GPC3 was expressed in Hep3B-SC22 cells and then the cells were fixed and stained with anti-β-Tubulin-AF647 antibody. B. The EGFP-GPC3 was expressed in Hep3B-SC22 cells and stained with the antibody against plasma membrane ganglioside G_M1_ that partitions into lipid raft (top row). The EGFP-Golgi was co-expressed mCherry-GPC3 in Hep3B-SC22 cells (middle row) and the mCherry-Calnexin was co-expressed with EGFP-GPC3 in Hep3B-SC22 cells (bottom row) for the identification of cytoplasmic GPC3 location. The cells were visualized with Nikon Sora Spinning Disk, 60x.

Previous research indicates that GPI-anchored proteins are associated with lipid rafts of cell membranes ^15^.To explore the membrane GPC3 distribution, lipid rafts on Hep3B cells were labeled and crosslinked by the antibody against plasma membrane ganglioside G_M1_, which selectively partitions into lipid rafts, and then the cells were visualized by confocal microscope. GPC3 was found to co-localize with lipid rafts on the Hep3B cell membrane, indicating the association with lipid rafts (Fig. 1B).

### GPC3 molecules are arranged into nano-clusters and the heparan sulfate chains promote cluster formation of Wnt3a-FZD1 on cell membrane

For many membrane proteins, molecular clustering plays a key role in their response to signaling. Thus, MINFLUX Nano-resolution microscopy was performed to obtain a molecular map of GPC3 on the cell membrane. The HS chains of GPI-anchored proteins are key factors in cell survival, Wnt3a signaling, maintaining the extracellular matrix (ECM) and mediating the interaction with heparin-binding proteins or ECM ^16,17^. To understand the role of HS chains in GPC3 spatial organization, we generated mCherry-GPC3 mutants with single and double HS chain binding site mutations (S495A and/or S509A) as illustrated (Fig.2A) and re-constituted the GPC3 proteins in Hep3B-SC22 cells. We confirmed the mutant phenotype by Western blots and flow cytometry. Western blots show decreased glycosylation of GPC3 in single-mutated constructs and complete loss of glycosylation for the double mutated construct, indicating successful mutation of the heparan sulfate binding sites (Fig. 2B). Further flow cytometry analysis showed that the depletion of HS chains does not decrease the translocation of GPC3 to the plasma membrane (Fig. 2C). In hepatoma carcinoma, the HS chains of GPC3 facilitate the Wnt3a signaling, and biologically, the loss of both chains results in significant decrease of Wnt3a reporter signaling (Figure 2D), indicating a critical role in signaling transduction.

**Figure 2.**
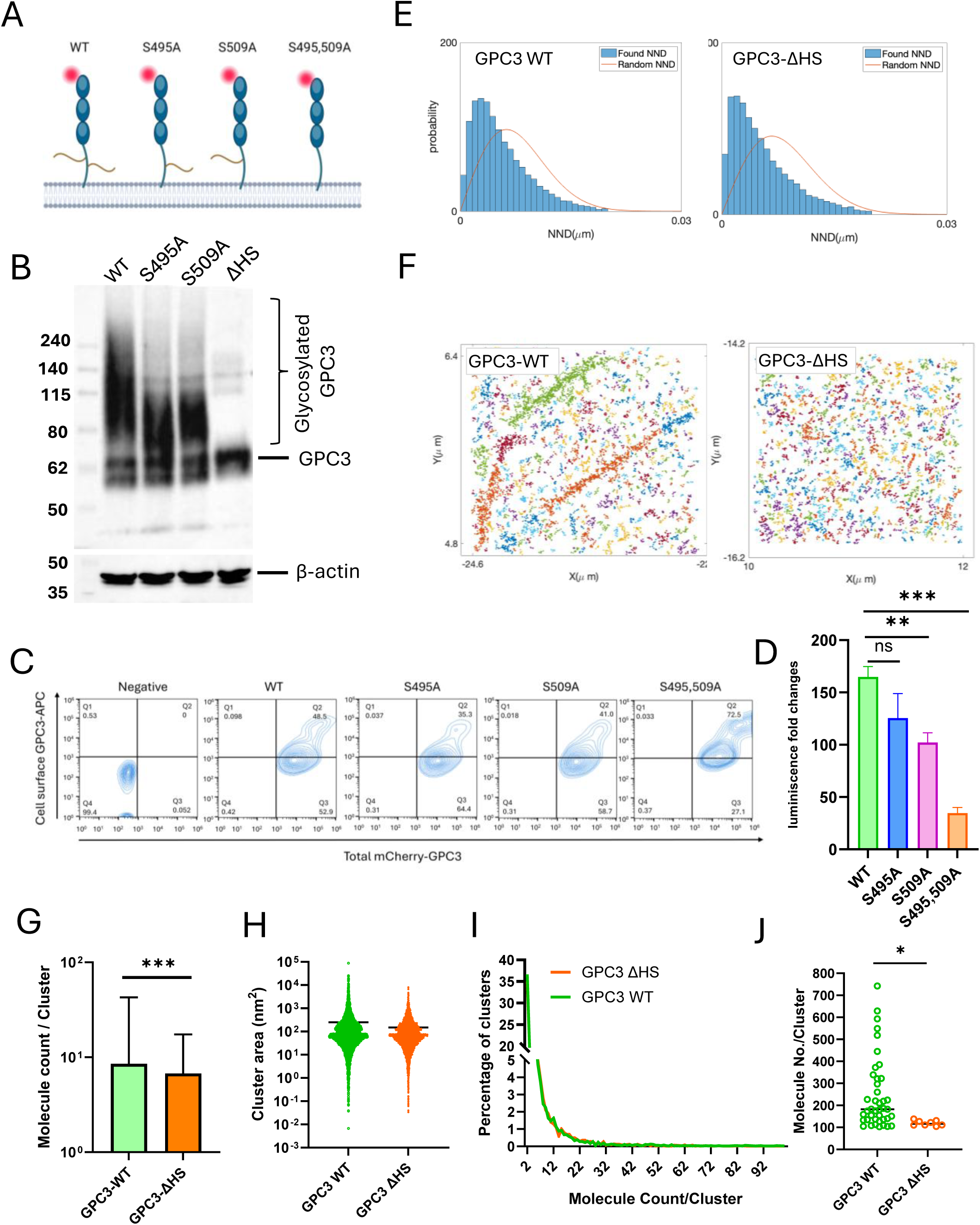
HS chains promote the formation of large GPC3 clusters. A. The illustration of GPC3 WT and mutants of S495A, 509A, and ΔHS. B. The depletion of HS chains eliminates glycosylation of GPC3. The GPC3 WT, S495A, S509A, and ΔHS constructs were expressed in Hep3B-SC22 cells, and then the cells lysates were cleared and precipitated with YP7 antibody against GPC3, and then the precipitated products were subjected to western blot detection for GPC3. C. Flow cytometry detection of membrane translocation rate of GPC3 mutants. mCherry-GPC3 WT and the three mutants were expressed in 293T cells, and then the cells were stained with mouse mCherry antibody and Goat anti-mouse secondary antibody conjugated with APC. The cells were subjected to spectral analysis by Sony ID7000 Analyzer. D. Wnt3a reporter assay with different GPC3 constructs. GPC3 WT and mutants of S495A, 509A, and ΔHS were transfected into HEK293-topflash cell line for 36 hours, then the cells were treated by Wnt3a conditioned medium for 6 hours, and the luminescence was read after cell lysis. Unpaired t test comparison was used to determine statistical significance, ***P < 0.005, ns: not significant. E. The proof of nonrandom distribution of GPC3 for WT and ΔHS constructs in Hep3B-SC22 cells. mCherry-GPC3 and ΔHS were expressed in Hep3B-SC22 cells and then the cells were stained with YP7-FLUX647 for MINFLUX Nano-resolution microscopy. The collected localizations were analyzed using a density-based clustering algorithm with a threshold distance of 20 nm to group the localizations associated to individual molecules. We then found the nearest neighbor distances between these molecules as shown by the histogram and compared to the distribution expected for a randomly distributed set of molecules with the same density (red curves). F. Rendered images for GPC3 clusters on Hep3B-SC22 cell membrane by MINFLUX Nano-resolution microscopy. G. Average GPC3 molecule count in the WT and ΔHS groups. H. Cluster area for the clusters of GPC3 WT and ΔHS. I. Proportion for GPC3 clusters with molecules count less than 100. J. Molecule count in the clusters that contain more than 100 molecules.

To compare the nanoscale distribution of GPC3 on cell membrane in the presence or absence of heparan sulfate chains, Hep3B-SC22 cells re-constituted with wild type GPC3 (mCherry-GPC3-WT) or GPC3 depleted of both HS chains (S495,509A/mCherry-GPC3-ΔHS). The cells were labeled with GPC3 mouse monoclonal antibody (YP7) conjugated with FLUX647 chromophore and visualized with MINFLUX Nano-resolution microscope. We analyzed clustering of the molecules by first identifying the mean position of each molecule by calculating the average coordinates of associated trace. These single positions were then grouped together using a density-based clustering algorithm with a distance threshold of 20 nm. We first compared the GPC3-WT and GPC3-ΔHS using data sets each containing 160,000 localizations of GPC3 molecules. The DBSCAN analysis confirmed the presence of both GPC3 single molecules and clusters on the cell membrane for both groups (Fig. 2E). There are 51% of GPC3 population existing as single molecules for GPC3-WT and 53% of the molecules are single for GPC3-ΔHS. We obtained the distribution of the number of molecules belonging to a single cluster and calculated cluster area based on the convex hull surrounding all molecules within a cluster (Fig. 2F, G), and the effective size of this area (Fig. 2H). For the GPC3 WT group, the mean count of molecules per cluster is 9 with an area of 250 nm^2^. The mean effective radius of WT clusters is 6.3 nm, with a minimum size of 0.046 nm, and maximum size of 167 nm. To evaluate the role of HS chains in GPC3 spatial localization, we further analyzed the molecule distribution of GPC3-ΔHS protein. The mean count of GPC3-ΔHS molecules per cluster is 7, which is a decrease of 22.2% compared to WT, and the mean area of clusters is 150 nm^2^ (Fig. 2G, I). This indicates less clustering of GPC3 in the absence of HS chains. Interestingly, although GPC3-WT and GPC3-ΔHS exhibited comparable distributions of small clusters (<100 molecules) (Fig. 2I), GPC3-WT uniquely supported the formation of high-occupancy clusters (Fig. 2J). Among the clusters analyzed per group, the GPC3-WT group formed 42 clusters containing more than 100 molecules, whereas GPC3-ΔHS formed only 8 clusters. For those large clusters, the mean molecular count is 247 per cluster with a mean radius of 50 nm, while the average molecular count of GPC3-ΔHS is only 119 per cluster (Fig. 2J). These data indicate that GPC3-WT protein promotes the maturation or stabilization of large supramolecular assemblies, whereas GPC3-ΔHS is largely restricted to forming smaller, less complex clusters.

As shown in Fig. 2D, the loss of HS chains impairs Wnt3a signaling, therefore, we aimed to unveil whether HS chains affect the Wnt3a coupling to the cellular receptors. To visualize the Wnt3a receptor complex on the plasma membrane, we incubated Hep3B-FZD1-mCherry cells with EGFP-Wnt3a conditioned medium. Confocal microscopy showed the association of Wnt3a on the plasma membrane of both cells that express GPC3-WT or GPC3-ΔHS, in addition, we also observed internalized Wnt3a coupling with GPC3 or FZD1(Fig. S1). However, the cluster details on the plasma membrane cannot be seen. Thus, we explored the cluster formation of GPC3-Wnt3a-FZD1 by three-dimensional multichannel (3D)-MINFLUX DNA-PAINT (Figure 3). The Hep3B-SC22-FZD1-mCherry cell line was transfected with plasmids expressing EGFP-GPC3 or EGFP-GPC3-ΔHS, and then the cells were incubated with SNAP-Wnt3a. The SNAP-Wnt3a were labeled with rabbit-anti-SNAP polyclonal antibody, and then the samples were labeled with oligo conjugated anti-GFP, anti-rabbit, and anti-mCherry nanobodies for EGFP-GPC3, SNAP-Wnt3a, and FZD1-mCherry staining, respectively. MINFLUX imaging of the three channels was performed by adding and washing out the respective imager strands in a sequential order, and the 3D and 2D images were rendered (Fig. 3A &B). To establish a null model for spatial association, 100 datasets consisting of three randomly distributed proteins with densities similar to those of the experimental data were generated and analyzed to obtain the co-localization rate for the random data. The co-localization ratio observed in the GPC3-ΔHS condition fell within the 95% confidence interval derived from the randomized simulations, indicating the absence of specific co-localization. In contrast, GPC3-WT displayed significant spatial organization, forming discrete clusters with Wnt3a and FZD1 (Fig. 3C). Pairwise co-localization analysis revealed that both GPC3-WT and GPC3-ΔHS formed specific clusters with Wnt3a. GPC3-ΔHS also exhibited specific clustering with FZD1. Strikingly, FZD1 and Wnt3a showed significant co-clustering only in the presence of GPC3-WT, whereas their spatial association in the GPC3-ΔHS condition was indistinguishable from random distribution (Fig. 3D). Besides, we found that the presence of HS chains increased the proportion of GPC3, Wnt3a and FZD1 homo-clusters, however, the total protein levels of Wnt3a binding to the cell surface is not elevated, suggesting that HS chains facilitate their nanoscale enrichment at the plasma membrane (Fig. 3E&F). Together, these data indicate that HS chains on GPC3 are required to stabilize the spatial association between FZD1 and Wnt3a despite preserved Wnt3a-GPC3 binding, providing a mechanistic basis for the attenuation of downstream signaling observed upon HS loss.

**Figure 3.**
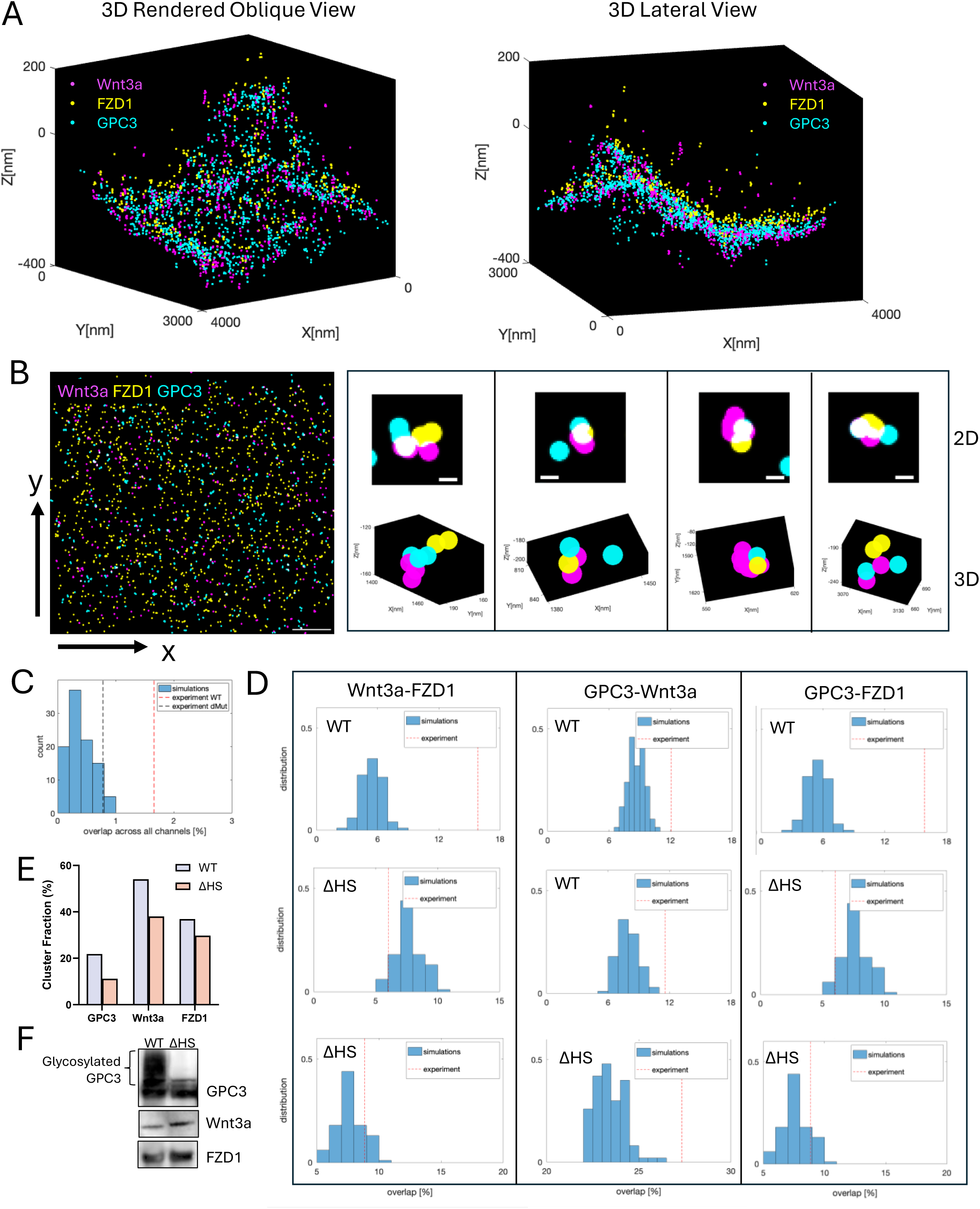
HS chains stabilize the Wnt3a receptor complex on the cell surface. A. 3D MINFLUX DNA-PAINT images of Wnt3a-GPC3-FZD1 complex at oblique view and lateral view. Hep3B-SC22-FZD1-mCherry cells were reconstituted with either EGFP-GPC3-WT or EGFP-GPC3-ΔHS for 36 hours, then the cells were incubated with SNAP-Wnt3a conditioned medium for 45min. After fixation, the cells were stained with anti-SNAP rabbit polyclonal antibody and subsequent nanobodies against GFP, mCherry, and rabbit antibody. MINFLUX imaging of the three channels were performed by adding and washing out the respective imager strands in a sequential order B. 2D rendered image of MINFLUX DNA-PAINT for Wnt3a-GPC3-FZD1 and the individual complex visualization at 2D and 3D mode, the top row are 4 respective complex in 2D mode, and the bottom row are the corresponding 3D images. Scale bar: 500 nm and 20 nm, respectively C. Data analysis of GPC3-FZD1-Wnt3a complex formation using nearest neighbor distances. The red dash line indicates the percentage of GPC3-WT complex formation, and the black dash line indicates the GPC3-ΔHS complex formation. The blue columns represent the range of non-specific coupling of random co-localization from the 100 repeats. D. Pairwise data analysis for the receptor couplings using the same method as panel C. E. Homo-cluster analysis for GPC3, Wnt3a, and FZD1, respectively. F. Protein level for membrane-bound Wnt3a in WT and ΔHS groups. Hep3B-SC22-FZD1-mCherry-EGFP-GPC3 WT/ΔHS stable cell lines were incubated with SNAP-Wnt3a conditioned supernatant for 40 min and then the cells were lysed after washing twice with 1xPBS for western blot analysis. GPC3 protein was probed by YP7 antibody, SNAP-Wnt3a was probed by anti-SNAP-antibody, and FZD1-mCherry was probed by anti-mCherry antibody.

### GPC3 is mobile and diffuses laterally on the hepatoma cell membrane

Membranous GPI-anchored proteins are dynamic in nature ^18^. To understand the dynamics of GPC3 associated with membranes in our hepatoma model, we analyzed dynamics of GPC3 in the membrane of Hep3B-SC22 cells that were re-constituted with mCherry-GPC3 by FRAP assay (50 repeats) (Figure 4A and Video 1). The intensity was reduced by photobleaching to a “baseline” (determined using fixed cells), and it then recovered as non-photobleached molecules moved into the photobleached area. The rate and extent of recovery depended on the contribution of different kinetic fractions. We analyzed the GPC3 molecule mobility and quantified fast-mobile, slow-mobile and immobile fractions (Fig. S1A-D). The fast-mobile fraction, F_0_, is estimated from the height of the first point in the curve; it contains molecules, which move so fast that they diffuse into the bleached area in the short interval between the end of photobleach and acquisition of the first post-bleach image frame. The slow-mobile fraction, F_S_, is estimated from the difference between the plateau in the curve and the initial point; the immobile fraction, F_imm_, defined as not recovering through the time of observation, is calculated as 1 – (F_0_ + F_S_).

**Figure 4.**
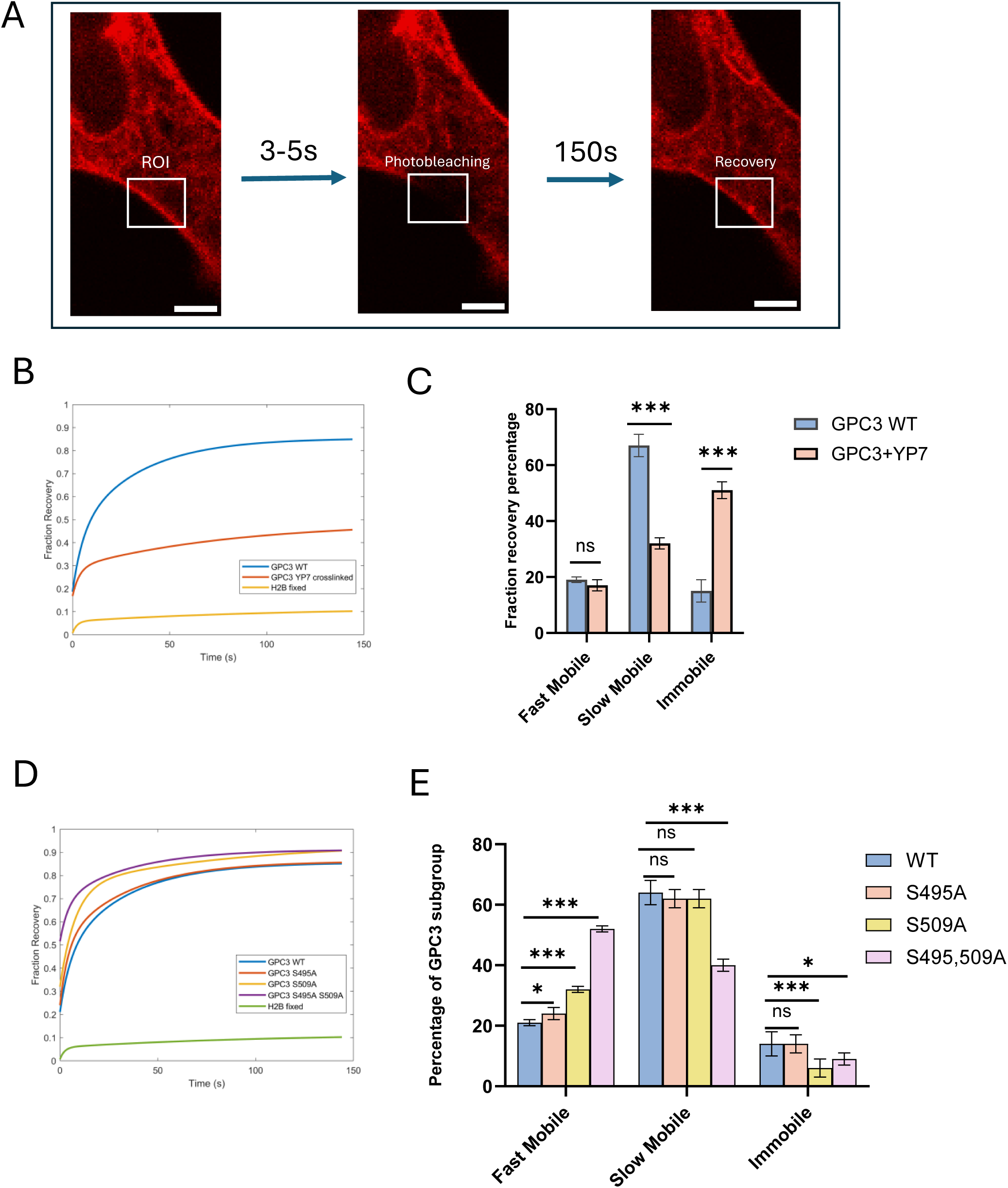
GPC3 is mobile on cell membrane and the deletion of heparan sulfate chains increases GPC3 mobility. A. Procedure for the mCherry-GPC3 bleaching on cell membrane. The Hep3B-SC22 cells were expressed with mCherry-GPC3 for 36h, and the membrane associated GPC3 in the ROI was photobleached and then the recovered fluorescence intensity in the ROI was recorded for 150s at a time interval of 0.48s. Scale bars: 3 µm. B. The recovery curve of mCherry-GPC3 in Hep3B-SC22 cells. The Hep3B-SC22 cells reconstituted with mCherry-GPC3 were incubated with mouse anti-GPC3 antibody YP7 and anti-mouse secondary antibody for crosslinking, and then the cells were subjected to FRAP assay. C. The fraction of GPC3 subgroup with different mobility with or without YP7 crosslink by FRAP. Data represent the mean ± SEM. Unpaired t test comparison was used to determine statistical significance, ***P < 0.001, ns: not significant, n ≥ 34. D. The recovery curve of WT GPC3 and three mutants by FRAP assay. The Hep3B-SC22 cells were transfected with mCherry-GPC3 WT, S495, S509, and ΔHS mutants for 36h, and then the cells were subjected to FRAP assay by Zeiss LSM780 microscope. E. The fraction of GPC3 subgroup with different mobility by FRAP. Data represent the mean ± SEM. Unpaired t test comparison was used to determine statistical significance, ***P < 0.001, ns: not significant, n = 50.

The average recovery half time for the slow-mobile fraction of WT GPC3 was 7.67 ± 1.34 s, accounting for a fraction of 67 ± 4%, with an immobile fraction of 14.6 ± 3.9%, and a fast-mobile fraction of 18.7 ± 1.5% (Fig. 4B and 4C, Table 1).

**Table 1.** Recovery half-life and immobile fraction of GPC3 and YP7 crosslinked GPC3 in FRAP.

|  | Mobile-Fast<br>Fraction | Mobile<br>Fraction | Mobile<br>Slow Recovery<br>time (s) | Slow<br>Half-Immobile<br>Fraction |
| --- | --- | --- | --- | --- |
| GPC3 WT | 0.19 ± 0.01 | 0.67 ± 0.04 | 7.7 ± 1.3 | 0.15 ± 0.04 |
| GPC3+YP7<br>crosslinking | 0.17 ± 0.02 | 0.32 ± 0.02 | 5.8 ± 1.8 | 0.51 ± 0.03 |
| H2B fixed | 0.01 ± 0.01 | 0.11 ± 0.02 | 3.4 ± 1.4 | 0.88 ± 0.02 |

Recovery after photobleaching may occur by exchange between the cytoplasmic pool of GPC3 and/or by lateral mobility of the GPC3 on the membrane. We immobilized GPC3 with antibody to determine the significance of lateral mobility during its recovery. We hypothesized that YP7 antibody crosslinking leads to immobilization of individual GPC3 by preventing lateral diffusion. We compared Hep3B-SC22 carrying GPC3 wild-type construct (50 repeats) and treated with YP7 antibody with non-treated control (34 repeats). Indeed, we observed that antibody treatment leads to a significant increase in the immobile fraction (14.6% ± 3.9%, versus 51.1% ± 2.7%).

The recovery rates were similar (their confidence intervals overlap: 7.66 ± 1.33 s versus 5.81 ± 1.75 s) as were the fast-mobile fractions (18.7 ± 1.5% versus 16.8 ± 1.7%) (Fig. 4B and 4C, Table 1). The result indicates that the recovery of GPC3 on the cell membrane is mainly due to lateral diffusion and not to the exchange between the cytoplasmic and membrane GPC3 pools. The antibody crosslinking could switch the slow-mobile GPC3 molecules to an immobile state but have no effect on the fast-mobile fraction.

### The heparan sulfate chains constrain GPC3 mobility

MINFLUX data reveal the effect of HS chains on GPC3 clustering. To evaluate whether HS chains restrict GPC3 mobility while mediating molecule coupling, we tested the effect of heparan chains on GPC3 membrane mobility by FRAP. We reconstituted mCherry-GPC3 mutants with single and double HS chain binding site mutations (S495A, S509A, or ΔHS) in Hep3B-SC22 cells for FRAP assay. For this experiment, we also repeated the FRAP of the wild type (50 repeats for the WT and three mutants each). The GPC3-ΔHS has a much larger fast-mobile population than the wild type (51.6 ± 1.1% versus 21.1 ± 1.4%) and the immobile fraction is smaller (8.8 ± 2.2% versus 14.4 ± 3.8%). For S509A, the immobile fraction is also smaller than the WT (5.7 ± 3.0% versus 14.4 ± 3.8%), while the fast-mobile group is slightly higher than WT (32.0 ± 1.4% versus 21.1 ± 1.4%) (Fig. 4D and 4E, Table 2). On the other hand, all parameters from S495A are very similar to the WT parameters. Furthermore, the inactivation of the S509 chain has a more significant effect on the GPC3 mobility than the S495 chain by FRAP. However, the mechanism by which WT GPC3 HS chains confine the mobility of GPC3 is yet to be discovered.

**Table 2.**
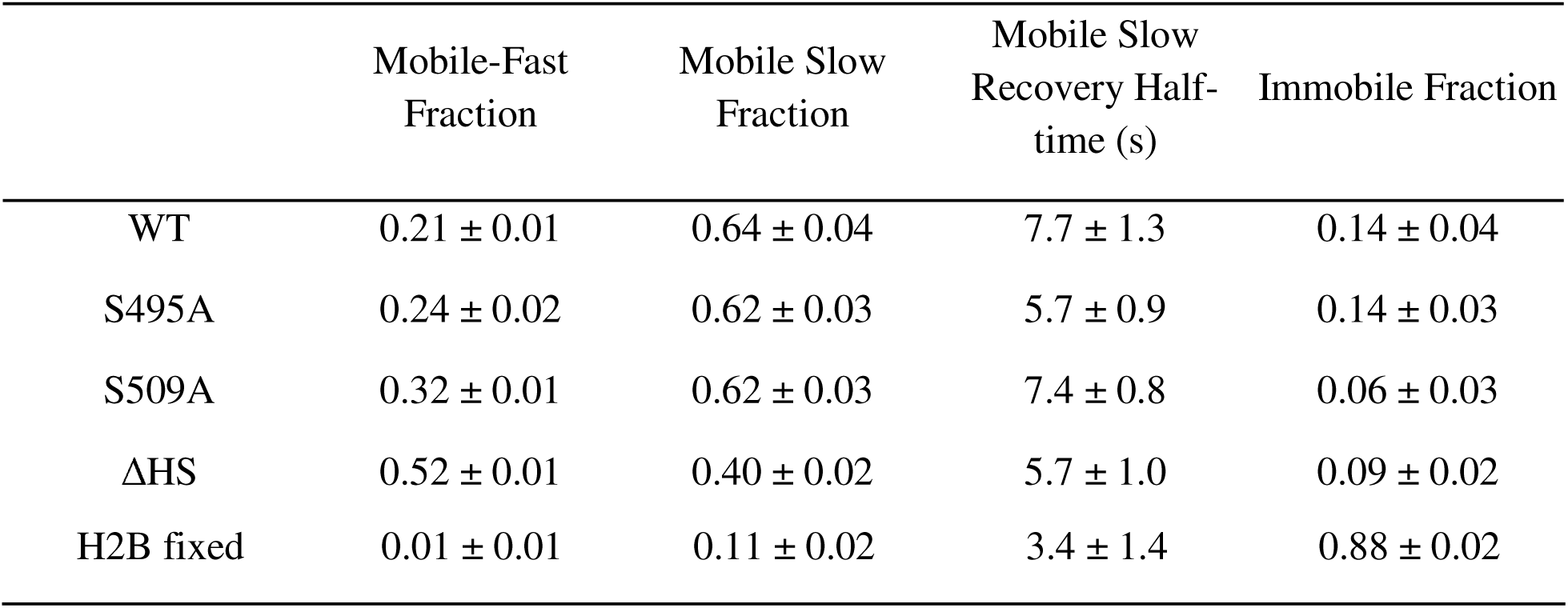
Summary for the mobility of WT GPC3 and mutants by FRAP assay.

### Single molecule tracking reveals that GPC3 diffused in a mixed pattern on the cell surface

As the FRAP data indicates that a fraction of GPC3 molecules are immobile, we decided to explore whether this is related to GPC3 clustering or corralling. We performed single molecule tracking for GPC3 on HepG2 cells with highly inclined thin illumination (HILO) microscopy. The GPC3 single molecules were labeled with 12pM YP7-Fab tagged with ATTO647N chromophore (Fab-ATTO647N), and the molecules were tracked at a time interval of 20ms for 24s (Video 2). Totally, 22,150 tracks were collected for GPC3 diffusion analysis. Individual trajectories of GPC3 molecules show that GPC3 undergoes mixed diffusion motion, with transition between Brownian and confined diffusion (Fig. 5A). We performed perturbation Expectation-Maximization (pEM) analysis to detect and characterize the states with molecular confinement, as described in Methods. In brief, we divided the tracks into smaller segments, 15 timepoints in length, to minimize the probability of transitions between states within a single track, and consequently 33,795 segments were obtained. For these segments, pEM revealed different states of confinement, five of which were observed with a frequency of ≥5% and thus were considered significant (Table 3). The significant states were characterized by their Mean Squared Displacement (MSD) calculated at various time-lags (Fig 5B). The time-dependent MSD curves of states exhibiting Brownian motion had a slope of ∼1 in log-log space, while slopes <1 are indicative of sub-diffusive motion and slopes >1 indicate super-diffusive, or directed, motion. We showed that 59.5% of the GPC3 population showed Brownian type motion without measurable confinement radii (states 4, 5, and 6). The remaining population was split among three sub-diffusive states: 4.1 % were confined within a radius of 50 ± 2 nm (state 1, D = 0.001 ± 0.0003 μm^2^/s), 14.1% within a radius of 113 ± 8 nm (state 2, D = 0.010 ± 0.002 μm^2^/s), and 14.7% within a radius of 185 ± 12 nm (state 3, D = 0.044 ± 0.008μm^2^/s) (Fig. 5C). Therefore, GPC3 membrane-associated molecules may transition between Brownian motion and restricted (confined) sub-diffusion. Confined sub-diffusive population may be further classified into the three states, differing by the radius of confinement and diffusion coefficients.

**Figure 5.**
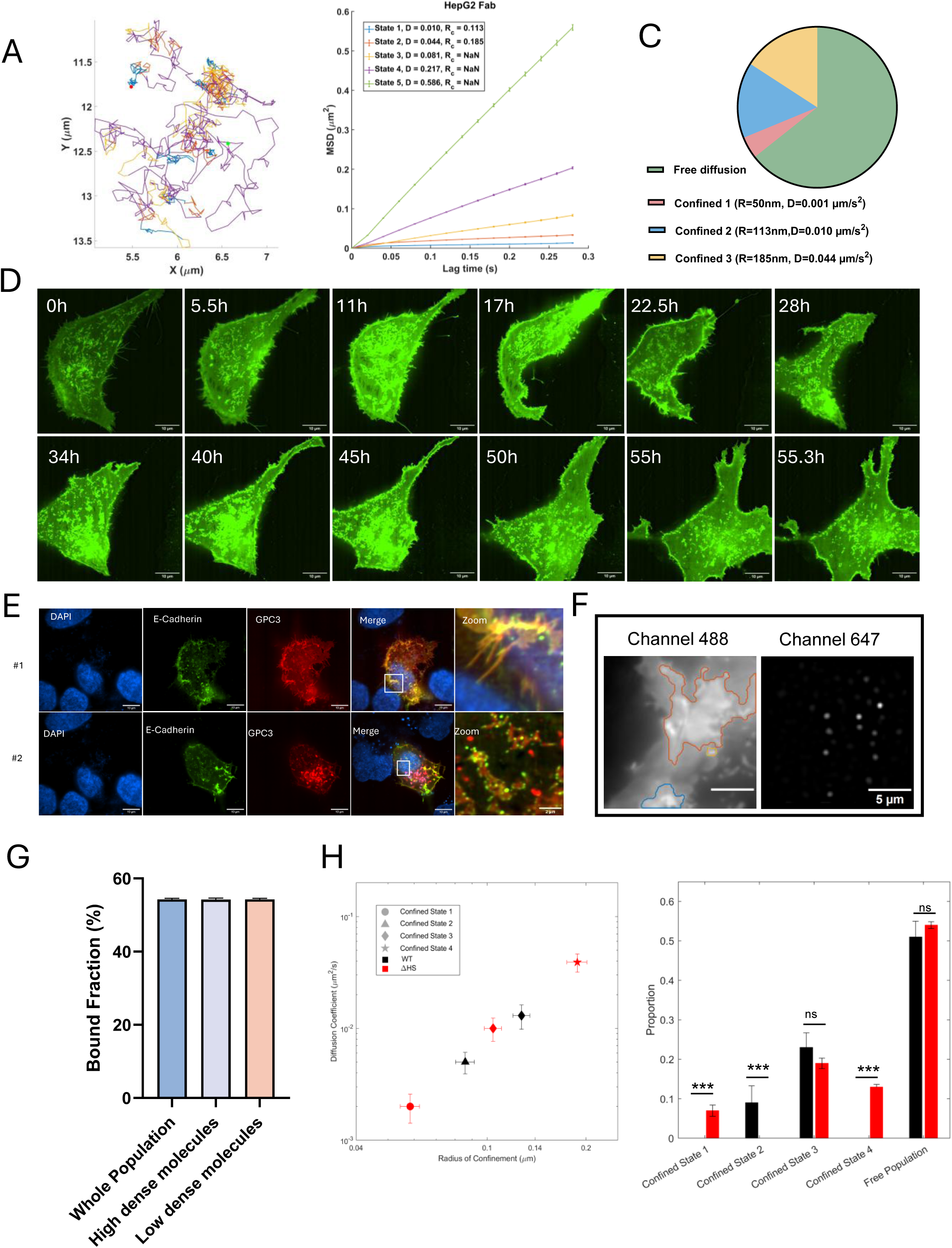
The loss of HS chains disrupts GPC3 diffusion at single molecule level. A. GPC3 diffusion trajectories on HepG2 cell membrane. The HepG2 cells were stained with YP7-Fab-ATTO647N at 12pM and subjected to HILO microscopy. The trajectories of GPC3 single molecules were plotted with a tracking duration of 23.7s. B. pEM analysis of GPC3 diffusion pattern on HepG2 cells. C. The fraction percentage for GPC3 in the states of Brownian diffusion and confined diffusion. D. Uneven distribution of GPC3 on the cell surface. The EGFP-GPC3 is expressed in Hep3B-SC22 cells for 36h, and then the cells were visualized by Nikon Sora confocal microscopy. Scale bars: 10µm. E. Confocal Microscopy for E-Cadherin and GPC3. GFP-E-Cadherin and mCherry-GPC3 were co-expressed in Hep3B-SC22 cells and then the imaged with Nikon Sora Spinning Disk Microscopy at an objective of 63x. #1 and #2 denotes two different cells. Scale bars: 10µm and 2 µm. (Inset magnification). F. Identification of high dense molecular raft by auto-thresholding of channel 488. The corresponding YP7-Fab-ATTO647N labeled GPC3 single molecules were present to the right. Scale bars: 2 µm. G. Bound fraction of GPC3 in the high dense molecular raft and low dense area. The GPC3 molecules were analyzed with residence analysis and compared in high dense molecular and low dense molecular area. H. The diffusion co-efficiency of GPC3 and proportion ofGPC3 diffusion states in GPC3 WT and ΔHS groups. The Hep3B-SC22 cells were transfected with EGFP-GPC3 WT or ΔHS for 36h, and then the cells were stained with YP7-Fab-ATTO647N for single molecule tracking by HILO microscope.

**Table 3.** pEM analysis of GPC3 molecules on HepG2 cells.

| State | Alpha | D ($\mu\text{m}^2/\text{s}$) | Rc (nm) | Proportion |
| --- | --- | --- | --- | --- |
| 1 | 0.32 | 0.001 | 50 | 4.1% |
| 2 | 0.45 | 0.010 | 113 | 14.1% |
| 3 | 0.5 | 0.044 | 185 | 14.7% |
| 4 | 1.02 | 0.081 | NaN | 10.4% |
| 5 | 1.03 | 0.217 | NaN | 25.5% |
| 6 | 1.06 | 0.586 | NaN | 23.6% |

### GPC3 exhibits the same mobility on the membrane in micro-domains regardless of the molecular density

From the confocal microscopy results, large bundles and clustering of GPC3 labeled with EGFP with relatively higher fluorescence intensity were observed at the cell surface in the movies obtained (Fig. 5D). The short bundles keep associating and dissociating with each other through a 55-hour time lapse duration of recording (Video 3). To further identify the potential structure that GPC3 associates with, we expressed the markers of focal adhesion, tight junction, adhesion junction, and desmosome junction. The result shows that the adhesion junction marker, E-cadherin, distributes along the GPC3 bundles on the cell membrane (Figure 5E), and we did not find proximal localization of GPC3 with other structures (data not shown), indicating that GPC3 may distribute in the same structure as adhesion molecules.

To identify whether the gathering of GPC3 is due to the sticking of immobile structures or corralling divided by micro-domains, we expressed EGFP-GPC3 in Hep3B-SC22 cells and gated the GPC3 molecules on 488nm channel by intensity thresholding (Fig. 5F). This allowed us to assay the dynamics of the molecule subpopulations in high dense areas and low dense areas separately (Video 4). We then analyzed and compared the molecular behavior between the different areas by gating based on the threshold. However, we did not find any difference for the GPC3 molecule residence time between the high dense areas and low dense areas (Fig. 5G), indicating the gathering of GPC3 is due to the corralling of molecules but not sticking to immobile structures. To further validate the speculation, we crosslinked GPC3 molecules with YP7 full antibody at the same molar concentration and found that in high dense area, GPC3 is easier to crosslink. In addition, there is a higher bound fraction in high dense area than low dense area, indicating that the higher dense area is formed by corralling but not interaction with a certain structure (71.4 ± 0.8% versus 66.2 ± 0.7%) (Fig. S2). The confined radius fell in the GPC3 cluster range in MINFLUX single molecule localization, indicating that the GPC3 clusters may correlate with the molecule corralling on the membrane.

### Removal of heparan sulfate chains results in a more heterogenous mobile behavior of GPC3

MINFLUX nanoscopy shows that GPC3-ΔHS forms similar clusters as GPC3-WT on the cell membrane when the clustered molecule is less than 100, but fewer clusters when the clustered molecules are more than 100. To understand the dynamic behavior of the molecules of GPC3-WT and GPC3-ΔHS, we reconstituted EGFP-GPC3-WT and EGFP-GPC3-ΔHS in Hep3B-SC22 cells and then analyzed the single molecule diffusion by HILO SMT microscopy and compared the molecular mobility with pEM analysis. The results show that for WT GPC3, 9.0% of the state exhibits confined diffusion with a radius of 86 ± 6 nm (state 2, D = 0.005 ± 0.001 μm^2^/s), and 23.4% with a radius of 127 ± 8 nm (state 3, D = 0.013 ± 0.003 μm^2^/s), accounting for 32.4% of the population (Table 4). There are 51% of GPC3 molecules undergoes Brownian diffusion state (states 4, 5, and 6) with higher diffusion coefficients (D=0.117-0.903 μm^2^/s).

**Table 4.** pEM analysis of EGFP-GPC3 molecules on Hep3B-SC22 cells.

| State | Alpha | D ($\mu\text{m}^2/\text{s}$) | Rc (nm) | Proportion |
| --- | --- | --- | --- | --- |
| 1 | 0.44 | 0.005 | 86 | 9.0% |
| 2 | 0.39 | 0.013 | 127 | 23.4% |
| 3 | 0.90 | 0.117 | NaN | 13.1% |
| 4 | 1.02 | 0.406 | NaN | 28.1% |
| 5 | 1.02 | 0.903 | NaN | 9.5% |

For GPC3-ΔHS, six diffusive states were resolved (Table 5). Three states displayed defined confinement radii (Rc = 58–188 nm) with diffusion coefficients ranging from 0.002 to 0.039 µm²/s, accounting for 39.2% of the population (Table 5). While GPC3-WT displayed distinct low-diffusion confined states with well-defined confinement radii, loss of heparan sulfate chains resulted in a redistribution toward faster, freely diffusing states, fewer deeply immobilized populations (state 1 in GPC3-WT), and more heterogeneous confinement. The remaining three states in the GPC3-ΔHS group exhibited Brownian diffusion with a higher diffusion coefficient (D = 0.079–0.746 µm²/s), together comprising 54.1% of the population. For the Brownian diffused molecules, the diffusion co-efficient of GPC3-WT concentrates more in the intermediate bin (0.406 µm²/s, 28.1%), whereas GPC3-ΔHS is more heterogenous and spread between low and high within the free pool. The presence of a distinct high-diffusion co-efficient state 6 in GPC3-ΔHS (22.1% at 0.746 µm²/s) suggests a subset of GPC3-ΔHS molecules experience much less hindrance, indicating that they diffuse faster, or they have fewer protein-protein contacts during the diffusion.

**Table 5.** pEM analysis for EGFP-GPC3-ΔHS molecule diffusion pattern.

| | Alpha | D ($\mu\text{m}^2/\text{s}$) | Rc (nm) | Proportion |
| --- | --- | --- | --- | --- |
| 1 | 0.36 | 0.002 | 58 | 6.9% |
| 2 | 0.40 | 0.010 | 104 | 18.9% |
| 3 | 0.49 | 0.039 | 188 | 13.4% |
| 4 | 1.05 | 0.079 | NaN | 7.7% |
| 5 | 1.05 | 0.254 | NaN | 24.3% |
| 6 | 1.07 | 0.746 | NaN | 22.1% |

Collectively, the deletion of HS chains does not dramatically change the overall proportion of Brownian versus confined diffusion states. Instead, HS loss redistributes GPC3 molecules across a broader range of diffusion coefficients and confinement radii, increasing diffusive heterogeneity and expanding the fast-diffusing population. Together with a redistribution toward higher-mobility SMT states, it suggests impaired persistence of confined assemblies. These findings indicate that HS chains are not required for initial membrane association but are critical for stabilizing GPC3 confinement within membrane nanodomains. This is consistent with the loss of large clusters in GPC3-ΔHS, a role for HS in promoting sustained molecular interactions and higher-order assembly. Despite this change, GPC3 retains the ability to associate with Wnt3a, as shown in MINFLUX DNA-PAINT, indicating that HS chains are not essential for direct ligand binding. Instead, HS chains appear to play a critical role in organizing the higher-order receptor assembly required for efficient FZD1-Wnt3a complex formation.

## Discussion

GPI-anchored proteins have been reported to be present on the cell membrane as nanoscale clusters and the organization of the GPI-anchored proteins are affected by the modulation of cholesterol and actin ^19–21^. In our confocal microscopy of GPC3, we found that GPC3 forms short bundles on the bottom membrane, and that these short bundles could concentrate during cell migration, resulting in rafts with high fluorescence intensity on the cell membrane. Further confocal microscopy shows that GPC3 localizes to regions enriched in the adhesion molecule E-cadherin (Fig.5E). After analyzing some high fluorescence areas by SMT, we found that they exhibit the same mobility as other areas at the single molecule level, indicating that they are correlated with certain structures but not sticking to them. It will be of interest to determine whether the corralling of GPC3 within membrane microdomains is mediated by adhesion-associated structures.

The HS backbone contains 50-400 monosaccharide units, which could be up to 200 nm in length. HS chains are involved in the cell survival signaling ^17^, motility ^7^ and contribute to the structural integrity of the ECM through interaction with laminin or collagens ^22,23^. In MINFLUX molecular mapping for GPC3, we observed the same distribution pattern for GPC3-WT and GPC3-ΔHS proteins for small clusters (containing less than 100 molecules). However, GPC3-WT contains more large clusters than GPC3-ΔHS, suggesting the HS chains facilitate forming higher-order assemblies. The larger clusters may imply the presence of multivalent interactions, stable nanoscale confinement, reduced lateral mobility, or longer residence times.

Consistent with the mapping result, the SMT shows both GPC3-WT and ΔHS proteins contain Brownian and confined diffused population, however, HS chains do not simply shift GPC3 from confined to Brownian motion but instead constrain the diffusion landscape, limiting heterogeneity and suppressing the emergence of highly mobile subpopulations, further supporting that HS chains function to stabilize and mature higher-order assemblies. In the FRAP assay, we also observed a much faster mobile fraction for GPC3-ΔHS protein.

In HCC model, upon Wnt3a binding, GPC3 forms a complex called the “signalosome” through association with Frizzled receptors, LRP5/6 coreceptors and cytoplasmic adaptors Dvl and Axin ^24^. Consistent with these observations, our super-resolution nanoscopy analysis revealed that GPC3-WT forms specific nanoclusters with Wnt3a and FZD1, whereas deletion of HS chains markedly disrupts this organization. Interestingly, even though we did not find less association of Wnt3a with plasma membrane, the reporter assay shows that there is no effective signal transduction without GPC3 HS chains, indicating GPC3 HS chains are required for the signal initiation regardless of Wnt3a binding (Fig.2D & Fig.3F). In addition, homo-cluster analysis revealed that GPC3, Wnt3a and FZD1 exhibited increased local concentration at the plasma membrane in the presence of intact HS chains. Pairwise clustering analysis further demonstrated that Wnt3a and FZD1 exhibit significant co-clustering only in the presence of intact HS chains, indicating that although Wnt3a associates with the plasma membrane, there is no efficient Wtn3a-FZD1 coupling (Fig.3D& E). The results further support a stabilizing role for GPC3 HS chains in receptor complex assembly.

Previous studies have shown that FZD1 can be degraded through ZNRF3-mediated ubiquitination; however, GPC3 HS chains facilitate the binding of R-spondin proteins to ZNRF3, promoting ZNRF3 degradation and thereby restoring FZD1 levels at the plasma membrane. Consistent with this mechanism, we observed fewer FZD1 molecules at the plasma membrane in the GPC3-ΔHS group after molecule normalization, suggesting enhanced FZD1 degradation in the absence of HS chains, which may be one of the mechanisms leading to the signaling attenuation.

These results support a model in which GPC3 HS chains act as nanoscale scaffolds that spatially organize and stabilize Wnt receptor complexes at the plasma membrane, thereby facilitating efficient signalosome formation and pathway activation (Figure 6).

**Figure 6.**
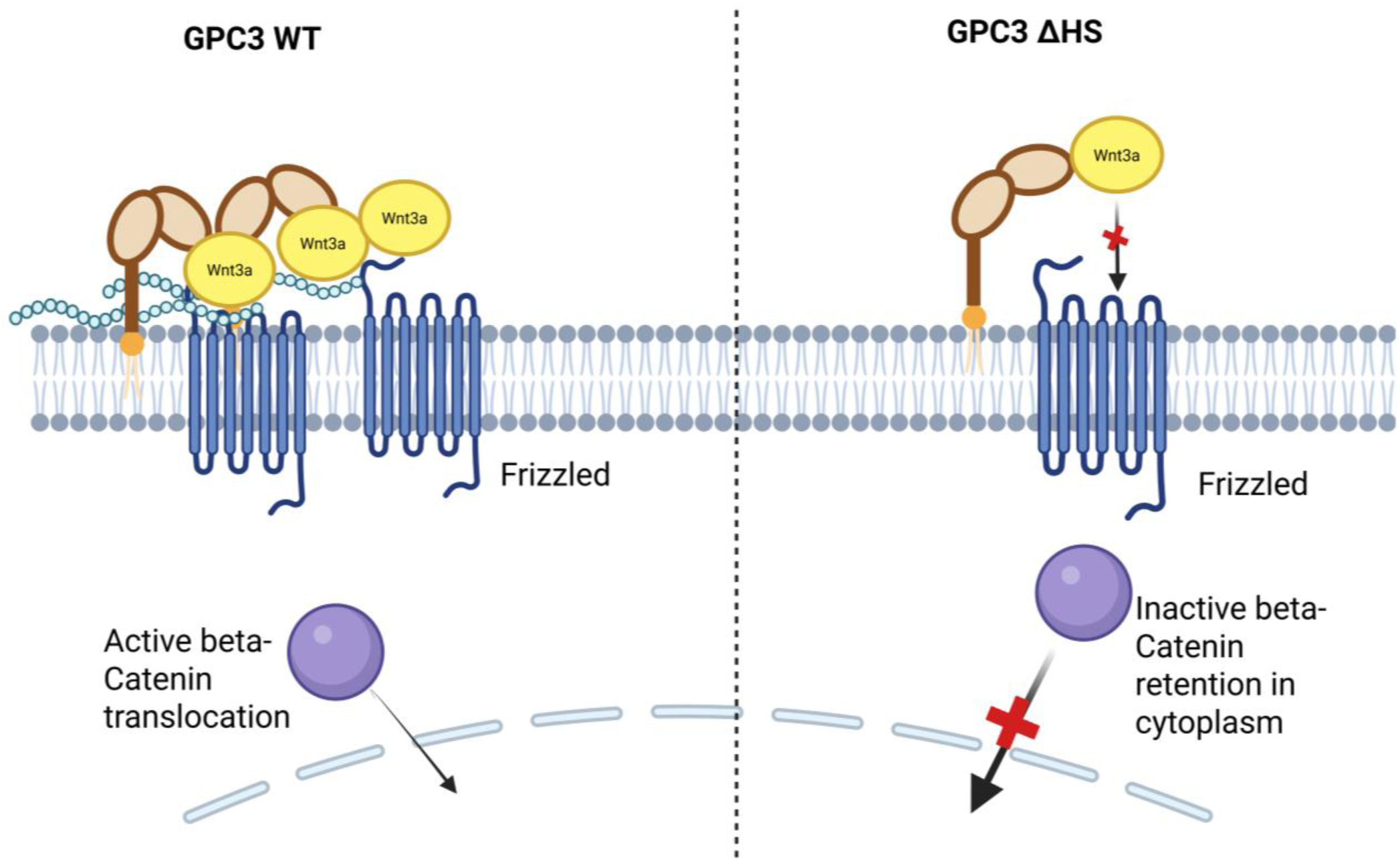
Cartoon illustration of the Wnt3a receptor binding mechanism. In the presence of GPC3 heparan sulfate chains, the Wnt3a could form complex with FZD1 and GPC3 receptors stably. However, when heparan sulfate chains are deprived, the coupling of Wnt3a and FZD1 is abolished while the Wnt3a-GPC3 coupling is not affected.

In conclusion, HS chains stabilize GPC3 on the HCC plasma membrane by promoting the formation and maturation of high-occupancy assemblies while preserving small-nanocluster formation. Loss of HS chains increases GPC3 mobility and heterogeneity, reduces deep confinement, and disrupts co-clustering with Wnt3a and Frizzled receptors. Together, these findings demonstrate that HS chains of GPC3 are critical for higher-order complex assembly and efficient Wnt3a signalosome formation in HCC models, thereby advancing our understanding of signaling mechanisms that drive cancer progression.

## Material and Methods

### Cell Culture and Medium

HepG2, Hep3B, and all the genetically engineered Hep3B cell lines were cultured using DMEM (Fisher Scientific, Cat# 11995081) supplemented with 1x Glutamax (Fisher Scientific, Cat# 35050079), 1x Penstrep (Fisher Scientific, Cat# 15140163), and 10% fetal bovine serum (GE, Cat# SH30071.03).

### DNA Constructs and Cell Transfection

GPC3 sequence (NM_001164618.2) and the mutants were cloned to PLM vector with an mCherry or EGFP tag inserted at the N terminus. EGFP-GalT plasmid is a gift from Jennifer Lippincott-Schwartz’s lab (Addgene plasmid # 11929, http://n2t.net/addgene:11929; RRID: Addgene_11929), mCherry-Calnexin-N-14 plasmid is a gift from Michael Davidson (Addgene plasmid # 55005; http://n2t.net/addgene:55005; RRID: Addgene_55005). EGFP-Wnt3a and FZD1-mCherry plasmids are gifts from Gary Davidson’s lab, Institute of Biological and Chemical Systems-Functional Molecular Systems (IBCS-FMS), Karlsruhe Institute of Technology (KIT), Karlsruhe, Germany. The plasmids were transfected to Hep3B-SC22 cells using jetOPTIMUS® DNA Transfection Reagent (Polyplus Transfection Inc, Cat# 101000051) according to the protocol.

### Immunoprecipitation and Western Blot

The Hep3B-SC22 cells were transfected with GPC3 constructs (WT, S495A, S509A, and ΔHS) for 36 hours, and then the cells were harvested by 1x RIPA buffer, with a protease inhibitor cocktail (Santa Cruz Biotechnology, CA, USA). The cell lysates were centrifuged at 12,000xg for 10 min to remove the cell debris, and the supernatant was incubated with humanized YP7 antibody for overnight at 4°C. 10 µl of supernatant was preserved for β-actin detection by western blot. The protein-antibody complex was then pulled down by KPL Protein A/G agarose (Seracare, MD, USA) at room temperature for 1 h. The complex was then resuspended in 1x RIPA buffer for western blot detection. The precipitated complex was separated in a 4-20% SDS-PAGE gel and transferred onto a nitrocellulose membrane. The membranes were probed overnight at 4°C using mouse YP7 primary antibody, followed by the horseradish peroxidase-conjugated secondary antibodies for 2 h at room temperature. Enhanced chemiluminescence (ECL) was used for the visualization of protein expression.

For Western blot of SNAP-Wnt3a and FZD1-mCherry proteins, the membrane transfer was done as above, and the proteins were probed with either SNAP antibody (Genscript Inc) or mCherry antibody (Takara Inc)

### Flow Cytometry

The expression level of reconstituted mCherry-GPC3 proteins (WT, S495A, S509A, and ΔHS) were determined on 293T cells by flow cytometry using 5 μg/mL of mCherry antibody (Invitrogen, MA, USA) and detected with a goat anti-rabbit IgG conjugated with Allophycocyanin (Jackson ImmunoResearch, Cat#711-116-152) at a dilution of 1:200. Data was visualized using SONY ID7000 (Sony Biotechnology) and analyzed by FlowJo v10.4.2 software (FlowJo, LLC).

### Confocal Microscopy

For confocal microscopy of fixed samples, Hep3B-SC22 cells were expressed with EGFP-GPC3 with the markers of Golgi or ER, and then the cells were fixed with 4% paraformaldehyde and 0.2% glutaraldehyde for 90 min for visualization with Nikon SoRa Spinning Disk Microscope, 60x Apo TIRF (N.A. 1.49) oil immersion objective and Hamamatsu ORCA Fusion BT sCMOS camera (Nikon Instruments Inc.).

For live cell tracking, Hep3B-SC22 cells were stably transduced with EGFP-GPC3 and were then tracked with Nikon SoRa Spinning Disk Microscope, 60x Apo TIRF (N.A. 1.49) oil immersion objective and Hamamatsu ORCA Fusion BT sCMOS camera (Nikon Instruments Inc.). All the images were denoised after acquisition using the Denoise.ai algorithm in the Nikon Elements software (V. 5.42).

### 3D MINFLUX DNA-PAINT

For 3D DNA-PAINT, SNAP-Wnt3a conditioned medium was incubated with Hep3B cells expressing EGFP-GPC3 for 1h at 37°C, and then the cells were fixed and stained with rabbit anti-SNAP primary antibody (Genscript Inc). Then the coverslip was washed and incubated with camelid nanobodies against GFP, mCherry, and rabbit antibody (Massive Photonics Inc). MINFLUX imaging of the three channels were performed by adding and washing out the respective imager strands in a sequential order.

The acquired datasets were first processed independently through the following steps: 1) finding the trace centers; 2) correcting drift using gold bead fiducial’s locations; 3) filtering out trace centers with precisions worse than 10 nm and traces containing fewer than four localizations; and 4) identifying and grouping trace centers corresponding to the same molecule. This workflow yields a set of molecule locations with high precisions for each dataset.

Next, the sequentially acquired data sets from GPC3, FZD1 and Wnt3a were registered using the same set of gold beads fiducials present across these datasets. Global shifts and rotations of the set of gold beads were determined with respect to a reference dataset (Wnt3a dataset) and used them to register all the molecule locations.

Finally, the reregistered molecule locations were used to quantify the ratio of overlapping molecules (co-localization) across each pair of channels, as well as across all three channels. To do so, we calculated nearest neighbor distances between molecules across sequentially acquired datasets, and molecules were considered overlapped if their separations were less than 3 times of the average precision (average precisions ∼6nm). For the sake of rigorous statistical comparison, we generated 100 simulated datasets with randomly distributed molecules at the same densities as the experimental data (i.e., the same number of molecules per unit area). Further, to assess overlapping ratios across all three channels (GPC3, FZD1 and Wnt3a), molecular densities were first uniformized for datasets acquired from WT and ΔHS samples. Finally, the overlapping ratios were considered statistically significant if the overlapping ratios from experimental data were larger than those obtained from randomly simulated molecules.

### Antibody or Fab Labeling

The YP7-Fab in PBS (in-house production, 1mg/ml) was adjusted to pH 8.0 and then incubated with ATTO647N dye (ATTO-TEC GmbH, AD 647) for 1h at room temperature, and then the labeled antibody was purified through PD-10 desalting columns packed with Sephadex G-25 resin (Cytiva, 17085101). For the labeling of the YP7 antibody with FLUX647 (Abberior, FX647), the antibody was modified with Traut’s reagent (2-imionothiolane in DMF), then the antibody purified with Sephadex G-25 resin. The purified antibody was incubated and purified as previously described. The labeled antibody was stored at 4 in PBS with 0.01% sodium azide.

### Wnt3a Luminescence Reporter Assay

The L-Wnt3a cells were cultured for 10 days, and the supernatant was collected on day 4 and 10 and combined as Wnt3a conditioned medium. The medium was stored at 4 °C. The cells were incubated with 50% Wnt3a conditioned medium for 6h and then lysed by passive lysis buffer (Promega, E194A). The lysed cells were incubated with Luciferase Assay Substrate (Promega, E151A) and the luminescence was read at the wavelength of 562 nm.

### Generation of L-SNAP-Wnt3a, L-EGFP-Wnt3a, Hep3B-SCC-FZD1-mCherry and Hep3B-SC22-FZD1-mCherry-EGFP-GPC3 stable cell lines

To generate the L-SNAP-Wnt3a or L-EGFP-Wnt3a stable cell line, we constructed a pLM-SNAP-Wnt3a and pLM-EGFP-Wnt3a plasmid in-house. Then the pLM-SNAP-Wnt3a or pLM-EGFP-Wnt3a plasmid was used for lentivirus production with the packaging plasmids, PAX2 and pMD2G in HEK-293T cells. The lentivirus was used to transduce the L cell line, and the positive clones were selected by limited dilution and verified by FACS using rabbit anti-SNAP antibody staining or GFP fluorescence. The cell supernatant was harvested 5 days after cell revival for incubation of the HCC cells.

To generate the Hep3B-SC22-FZD1-mCherry cell line, the FZD1-mCherry sequences were subcloned to pLM plasmid, and lentivirus was generated from the plasmid using the same packaging system as above for transduction of Hep3B-SC22 cells. The positive cells were sorted by flow cytometry (BD FACSAria Fusion).

To generate Hep3B cell line that express both FZD1-mCherry and EGFP-GPC3/ΔHS, Hep3B-SC22-FZD1-mCherry cell line was transduced with EGFP-GPC3 WT or ΔHS lentivirus, and the positive cells were selected by cell sorting (BD FACSAria Fusion).

### Fluorescence Recovery after Photobleaching (FRAP)

Wild type GPC3 and mutant GPC3 (S495A, S509A, ΔHS (S495&509A)) were expressed with mCherry tag in the Hep3B-SC22 cell line. The cells were imaged at 36 hours after transfection. Imaging conditions: FRAP was performed on LSM780 confocal microscope (Carl Zeiss, Inc) with 63x oil immersion objective. Cells were imaged with a 594 nm laser. A rectangular region was photobleached on the cell membrane with a short laser pulse operating at the maximum laser power. Fluorescent recovery was monitored at 0.48 sec time intervals. Image background was subtracted from those measurements. Intensity of the bleached area was divided by the intensity for the non-bleached membrane area in the same cell to correct for bleaching due to imaging. The resulting curve was normalized to the pre-bleach level of brightness. Normalized curves from the individual cells were averaged.

Ideally, photobleaching would result in the disappearance of all fluorescence within the measurement region, and the initial point in the FRAP curve would give information on the fraction of the fluorophores that recover between the end of the bleach and the first FRAP measurement. However, reversible photobleaching and background also contribute to the initial point, and we therefore used fixed H2B-mCherry to normalize the averaged FRAP curves. Re-normalized curves were then fit by a two-component exponential recovery. The fit parameters include the fractions of mobile-fast (F0), the two mobile-slow (F1, F2), and immobile molecules (Fimm), as well as the recovery rates of the two mobile-slow fractions (k1, k2). For simplicity, we report the total mobile-slow population (FS = F1 + F2) and the average half-time recovery 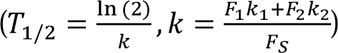 of the mobile-slow fractions along with the fractions of mobile-fast and the immobile populations for the different conditions and mutants as described in Kimura and Cook (2001) ^25^.

### Single Molecule Tracking

Sparce labeling in 647 nm channel was used to track individual GPC3 molecules on hepatoma cell lines. GPC3 was stained by 12pM YP7-Fab derived from YP7 mouse IgG antibody conjugated with ATTO647N chromophore. Stained cells were washed with PBS 4 times and then incubated in Fluorobrite DMEM for imaging recording.

The data were collected on a custom-built HILO-illumination microscope ^26^. The 488 nm laser line was used to illuminate Alexa488 whereas the 647 nm laser line was used to illuminate Atto647. Two channels were collected simultaneously. Time-lapse movies were acquired as 16-bit single focal plane images with 20 ms laser irradiation (Ex: 1 mW 488 nm; Ex: 3 mW 647nm) and 20 ms time interval for a total of 24 s.

We used the custom-made software TrackRecord ^27^ in MATLAB (The MathWorks, Inc., Natick, MA). To analyze each time series, data were filtered using top-hat, Wiener, and Gaussian filters. A region of interest (ROI) was defined to encompass the nuclei based on the nuclear GFP staining, then nuclear particles were detected, fitted to two-dimensional Gaussian function for subpixel localization, and finally tracked using a nearest neighbor algorithm ^28^. The tracking parameters were as follows: window size for particle detection 3 pixels, maximum frame to frame displacement of 6 pixels, shortest track 2 frames, and gaps to close 2.

The bound-fraction of each population was obtained by finding the ratio of immobile track segments to all track segments. Immobile segments of tracks were identified based on their displacements compared to HaloTag-H2B data, which was acquired with identical imaging conditions as GPC3. Because the motion of H2B includes the thermal motion of chromatin, it can be used to determine the maximum distance a DNA-bound molecule will be allowed to move. Therefore, we used the jump distribution of H2B tracks to define three parameters: Rmin, Rmax, and Nmin. Rmin is the maximum distance that can be moved between two consecutive frames and is the 99th percentile for H2B tracks. Similarly, Rmax is the 99th percentile of distance traveled by H2B at a time-lag equal to the shortest bound track, Nmin. Nmin is chosen to reduce the probability of a diffusive track as bound. For the data here, we used Rmin = 0.22 mm, Rmax = 0.315 mm, and Nmin = 4 frames.

Track segments were classified into distinct diffusive states using perturbation-Expectation Maximization^29^. Assumptions about the type of diffusion exhibited by the tracked particles are not required with pEM and the number of diffusive states can be deduced from the analysis. pEM analysis requires all analyzed tracks to be of the same length, and it is preferable to use shorter tracks that do not include transitions between states. Therefore, tracks were split into 15 frame (280 ms) segments and the pEM classification analysis was performed on the set of all these track segments. The minimum number of states for the system to converge to was set at 2 and the maximum at 10. If the optimal number of states that the analysis converged to was 10, the algorithm was rerun with a higher number of maximum states. The number of reinitializations was set to 50 with 200 perturbation trials. The maximum number of iterations was 1000 with a convergence criterion for the change of log-likelihood of 10–7. The number of features for the covariance matrix was set to 3 for tracks of length 7. A motion blur coefficient was calculated as (1/6) (Δ_e_/Δ_t_), where Δ_e_, corresponds to the exposure time and Δ_t,_ the features for the covariance matrix was set to 3 for tracks of length 7. A motion blur coefficient acquisition interval.

States obtained from pEM analysis that consisted of at least 5% of the total population were considered significant states and characterized further using the plots of their Mean Squared Displacements (MSD) over all time lags. The MSD(t) curve can be approximated as an exponential growth with rate constant, α.

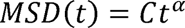

Brownian motion is characterized by α ≈ 1, while α < 1 indicates sub-diffusive motion, and α > 1 shows super-diffusive motion. In practice, noisy measurements can cause deviation from one for Brownian motion, and we therefore specify Brownian motion as having 0.7 ≥ α > 1.1. For states with α < 0.7, we fit the MSD curve to a confined diffusion model, 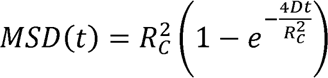 to obtain the radius of confinement, Rc.

### MINFLUX Nano-resolution Analysis

For MINFLUX nanoscopy analysis, images were acquired using a commercial MINFLUX system (Abberior, Gottingen, Germany). HepG2 cells were cultured on #1.5 18-mm round glass coverslips (Warner instruments, 64-0714) for 2 days. The cells were fixed with 4% paraformaldehyde and 0.2% glutaraldehyde for 90 min. After washing the cells 3 times with PBS, the cells were stained with YP7-FLUX647 antibody with 0.5% BSA for 50min at room temperature. The cells were washed with PBS 5 times and then preserved with 1xPBS for imaging.

Coverslips were mounted into the buffer with 18.8 mM beta-mercaptoethylamine and oxygen scavengers [50mM Tris–HCl (pH 8.0), 10mM NaCl, 10% (w/v) glucose, 64 μg/ml catalase (Sigma-Aldrich), 0.4 mg/ml glucose oxidase (Sigma-Aldrich)] to regulate photo blinking of FLUX647.

To correct for XYZ drift, 150 nm gold bead fiducials were applied to the coverslip prior to mounting (BBI solution, SKU EM. GC150). Samples were sealed with silicone glue and imaged.

2D MINFLUX measurements were performed with a 647 laser at 19 mW in the first iteration, and a pinhole size of 0.83 AU.

Data from MINFLUX consists of coordinates obtained by repeated localization for each single fluorescent molecule of fluorophores that are grouped together into traces. Any trace that contained less than 5 localizations was discarded to remove localizations due to noise. The remaining traces were then binned into pixels of 4 × 4 nm to generate a pseudo image where the brightness of each pixel represents the number of localizations in that area.

Tracks with at least 5 localizations were analyzed as clusters by first finding the average position of each trace. The average positions are considered the positions of individual molecules, which are then clustered using a density-based clustering algorithm (DBSCAN) ^30,31^ with a maximum distance of 20 nm between molecules.

### Statistical Analysis

The statistical analysis was done by GraphPad Prism 10 Software (GraphPad Software, Inc., San Diego, CA, USA). All data were conducted in triplicate and represented as means ± the standard error of the mean (SEM). The comparison between the two experimental groups was performed by Student’s unpaired t-test (two-tailed). Comparisons of the extracted FRAP populations and populations obtained from pEM analysis were performed using a z-test.

## Acknowledgements

We thank Dr. Michael J. Kruhlak for the microscope training, suggestions, and revisions on the manuscript writing. We thank NIH Fellows Editorial Board for editing and revising the manuscript. The contributions of the NIH author(s) are considered Works of the United States Government. The findings and conclusions presented in this paper are those of the author(s) and do not necessarily reflect the views of the NIH or the U.S. Department of Health and Human Services.

## Author contributions

Conception and design: S.L., M.H., and T.K.; Development of methodology: D.B., T.K., and F.M; Acquisition of data: S.L. and T.K.; Analysis and interpretation of data: S.L., T.K., D.B., F.M., and M.H.; Manuscript preparation: S.L., T.K., F.M., and D.B.; Revision and proofreading of manuscript: S.L., T.K., D.B., F.M., and M.H. All authors read and approved the final version.

---

**Figure S1.**
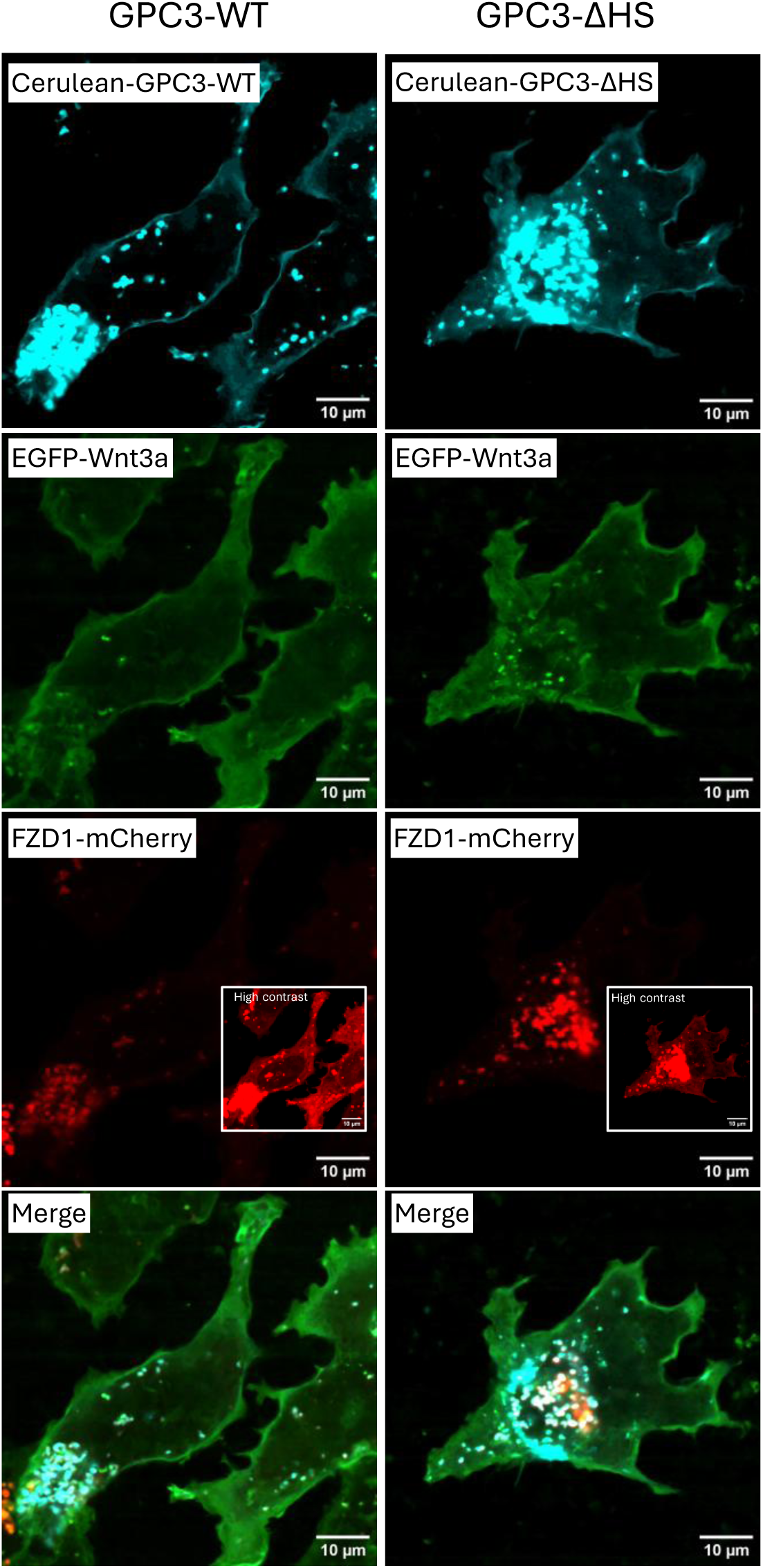
Confocal microscopy for Wnt3a binding visualization on HCC cell surface. Hep3B-SC22-FZD1-mCherry cells were transfected with Cerulean-GPC3-WT or ΔHS for 36 hours and then incubated with EGFP-Wnt3a conditioned supernatant for 40 min. The cells were visualized with Leica Stellaris 8 FLIM microscope, 63x.

**Figure S2.**
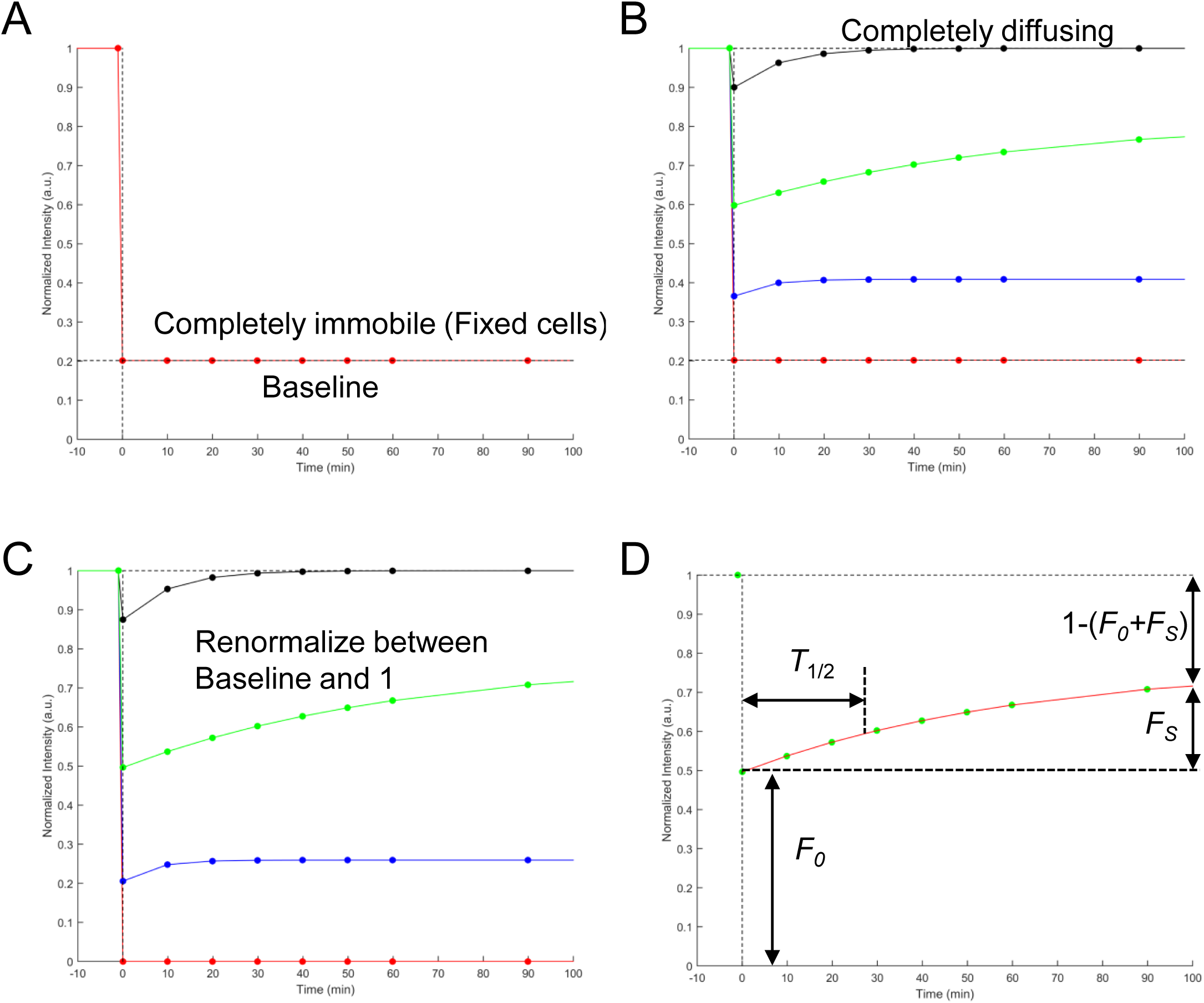
Schematic of FRAP renormalization and curve fitting. A. To extract quantitative values from FRAP curves, we must first measure the recovery curve for a completely immobile protein, such as H2B in fixed cells. The depth of the bleach is considered the Baseline with which other FRAP curves are compared against. B. Experimental FRAP curves will then lie between Baseline and 1, which represents full recovery. C. All curves are then renormalized, such that the Baseline becomes zero. D. Fitting the FRAP curve to an exponential recovery result in the parameters describing the recovery. These parameters consist of: the fast-mobile fraction, F_0_, which is estimated from the height of the first point in the curve; the slow-mobile fraction, F_S_, which is estimated from the difference between the plateau in the curve and the initial point; the immobile fraction, F_imm_, which is equal to 1 – (F_0_ + F_S_); and the half-time of recovery, T_1/2_.

**Figure S3.**
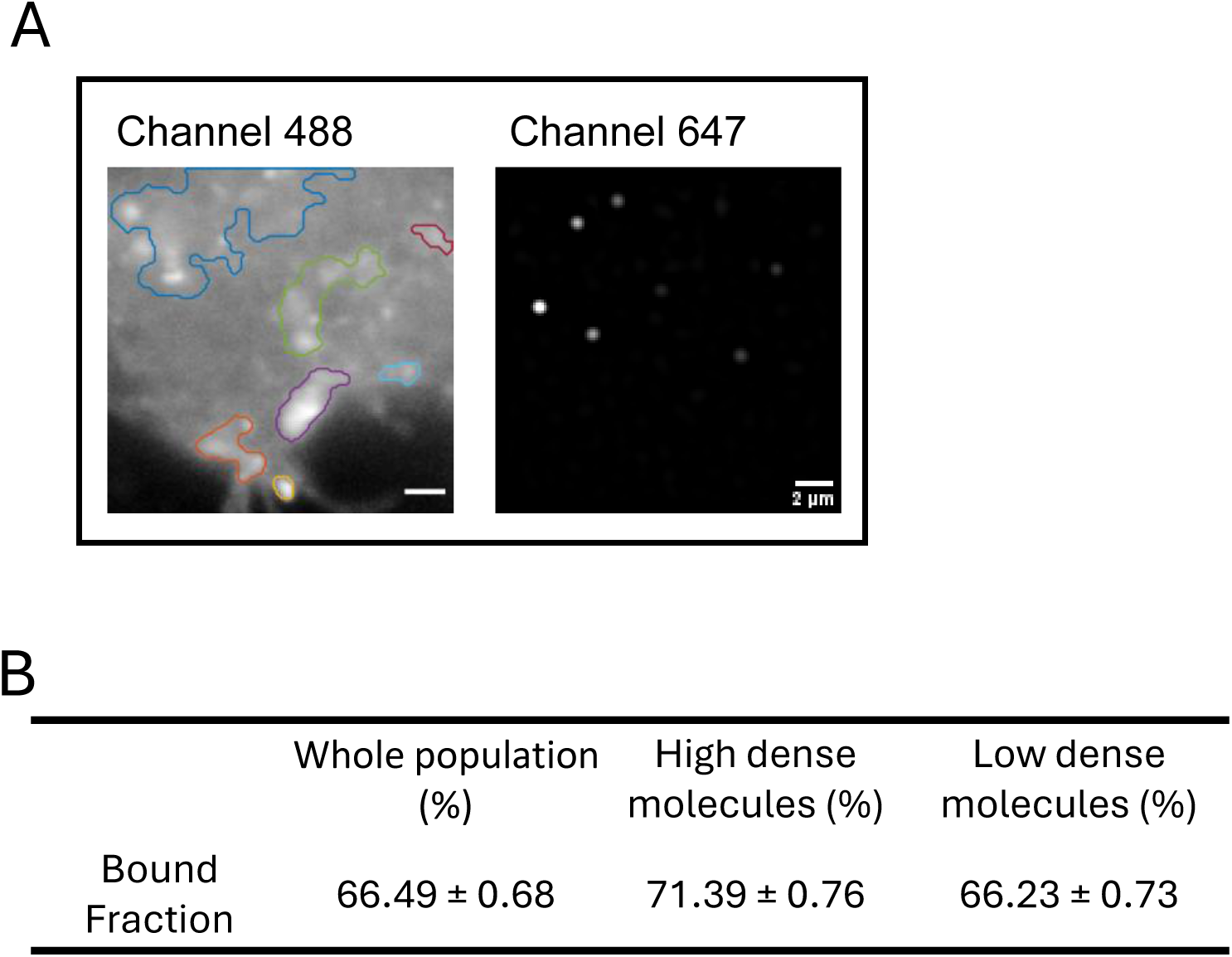
Comparison of YP7-labeled GPC3 on the cell membrane. A. Identification of high dense molecular raft by auto-thresholding of channel 488. The corresponding YP7-ATTO647N labeled GPC3 single molecules were present to the right. Scale bars: 2 µm. B. Bound fraction of GPC3 molecules in the high dense molecular raft and low dense area. The GPC3 molecules were analyzed with residence analysis and were compared in high dense molecular and low dense molecular area.

**Video 1. GPC3 is very mobile by FRAP assay.** The mCherry-GPC3 was re-constituted in Hep3B-GPC3-KO cells for 36 hours, and then the cells were subjected to FRAP assay by Zeiss LSM780 with a time interval of 0.48s. The duration of the video is 150 seconds.

**Video 2. GPC3 single molecule tracking on HepG2 cell surface.** The HepG2 cells were stained with 200pM YP7-ATTO647 for single molecule tracking by HILO microscopy. The time interval of the frames is 20ms with 1,200 frames per video.

**Video 3. GPC3 forms short bundles or concentrates in local domains on the cell membrane.** Confocal microscope tracking for GPC3 in Hep3B-SC22-EGFP-GPC3 stable cell line. The Hep3B-SC22-EGFP-GPC3 cells were cultured in chambers and subjected to live cell tracking by Nikon Sora Spinning disk Microscope at an objective of 60x. The time interval is 15min and the duration is 55 hours.

**Video 4. HILO SMT microscopy tracking for EGFP-GPC3 on Hep3B-SC22 cells.** The EGFP-GPC3 was expressed in Hep3B-SC22 cells and stained with 200pM YP7-ATTO647 for single molecule tracking by HILO microscope with channel 488 and 647. The time interval of the frames is 20ms and 1,200 frames were collected per video.

**Video 1. GPC3 is very mobile by FRAP assay.** The mCherry-GPC3 was reconstituted in Hep3B-GPC3-KO cells for 36 hours, and then the cells were subjected to FRAP assay by Zeiss LSM780 with a time interval of 0.48s. The duration of the video is 150 second.

**Video 2. GPC3 single molecule tracking on HepG2 cell surface.** The HepG2 cells were stained with 200pM YP7-ATTO647 for single molecule tracking by HILO microscopy. The time interval of the frames is 20ms with 1,200 frames per video.

**Video 3. GPC3 forms short bundles or concentrates in local domains on the cell membrane.** Confocal microscope tracking for GPC3 in Hep3B-SC22-EGFP-GPC3 stable cell line. The Hep3B-SC22-EGFP-GPC3 cells were cultured in chambers and subjected to live cell tracking by Nikon Sora Spinning disk Microscope at an objective of 60x. The time interval is 15min and the duration is 55 hours.

**Video 4. HILO SMT microscopy tracking for EGFP-GPC3 on Hep3B-SC22 cells.** The EGFP-GPC3 was expressed in Hep3B-SC22 cells and stained with 200pM YP7-ATTO647 for single molecule tracking by HILO microscope with channel 488 and 647. The time interval of the frames is 20ms and 1,200 frames were collected per video.

